# Nondisjunction and unequal spindle organization accompany the drive of *Aegilops speltoides* B chromosomes

**DOI:** 10.1101/547612

**Authors:** DanDan Wu, Alevtina Ruban, Jörg Fuchs, Jiri Macas, Petr Novák, Magdalena Vaio, YongHong Zhou, Andreas Houben

## Abstract

Supernumerary B chromosomes (Bs), which are often preferentially inherited, deviating from usual Mendelian segregation. This chromosome drive is one of the most important features of Bs. Here we analyzed the drive mechanism of *Aegilops speltoides* Bs and provide direct insight into its cellular mechanism. Comparative genomics resulted in the identification of the tandem repeat AesTR-183 of *Ae. speltoides* Bs, which also can be found on the Bs of *Ae. mutica* and rye, was used to track Bs during microgametogenesis. Nondisjunction of CENH3-positive, tubulin interacting B sister chromatids and an asymmetric spindle during first pollen grain mitosis are likely components of the accumulation process. A quantitative flow cytometric approach revealed, that independent on the number of Bs present in the mother plant Bs accumulate in the generative nuclei with more than 93%. Nine of eleven tested (peri)centromeric repeats were shared by A and B chromosomes. A common origin of the drive process in *Poaceae* is likely.

## Introduction

Supernumerary B chromosomes (Bs) are considered as dispensable chromosomes, which under standard conditions do not confer advantages on the organisms that harbour them. Bs are found in thousands of fungi, plant and animal species and are thought to stand for a specific type of selfish DNA (Jones and Rees, 1982). Recent advances in deciphering the sequence composition of B chromosomes provided a unique opportunity for the analysis of the origin and evolution of this enigmatic genome component. It was shown that the Bs of several plant and animal species harbour gene-derived sequences. This allowed to trace their origin from duplicated fragments of multiple A chromosomes of their host species (reviewed in (Ruban et al., 2017). Hence, Bs could be considered as a by-product of standard chromosome (As) evolution.

Often the transmission rate of B chromosomes is higher than 0.5, not obeying the Mendelian law of equal segregation. The resulting transmission advantage is collectively referred to as ‘drive’. The maximum number of Bs tolerated by the host varies between species and depends on a balance between the B chromosome drive, and the negative effects, especially on fertility and vigour, caused by B chromosomes if they occur in higher numbers (Bougourd and Jones, 1997). Among the relatively small number of B chromosome-positive species that have been investigated, relative to the total number of known species, there is a variety of mechanisms of B chromosome drive that occurs before, during or after meiosis, while in some cases drive has not been found (reviewed in (Austin et al., 2009; Burt and Trivers, 2006; Carlson, 1988; Houben, 2017; Jones, 1991; Jones, 2018)). Besides drive, the non-Mendelian inheritance of Bs could also be affected by mitotic and meiotic instabilities.

The drive of the rye (*Secale cereale* L.) B chromosome is one of the best-analyzed accumulation mechanisms amongst supernumerary chromosomes. Rye Bs do not separate at anaphase of the first pollen grain mitosis and mostly are included in the generative nucleus. In the second pollen mitosis, the Bs segregate normally into both sperm nuclei (Hasegawa, 1934). Nondisjunction of rye Bs is likely caused by extended cohesion of the B sister pericentromeres. Because of unequal spindle formation at the first pollen mitosis, nondisjoined B chromatids preferentially become located toward the generative pole and get consequently included into the generative nucleus. The failure to resolve pericentromeric cohesion is under the control of the B-specific nondisjunction control region (reviewed in (Houben, 2017)). A comparable accumulation mechanism takes place in the female gametophytes of rye (Hakansson, 1948).

In *Aegilops speltoides* Tausch (syn. *Triticum speltoides* (Tausch) Gren.), an annual diploid Poaceae (genome type: S), besides the seven pairs of A chromosomes, plants with a maximum number of eight additional Bs were reported (Raskina et al., 2011). The particular number of Bs is constant in all aerial parts within the same individual. Interestingly, B chromosomes are completely absent in the roots (Mendelson and Zohary, 1972; Ruban et al., 2014). Based on FISH experiments using *Spelt I* and 5S rDNA probes the B of *Ae. speltoides* is considered to be derived from the A chromosomes 1, 4 and 5 (Belyayev and Raskina, 2013; Friebe et al., 1995; Raskina et al., 2011).

Reciprocal crosses between 0B and +B plants indicated that the drive of Bs occurs during the first mitosis in the male gametophyte while there is no accumulation of Bs during the development of female gametophytes (Mendelson and Zohary, 1972). The non-Mendelian transmission of Bs cannot be solely explained by irregular segregation during meiosis. The net increase of B number in the progeny of 0B × 1B crosses can only be accounted for the directed nondisjunction of the Bs to the generative nucleus during first pollen mitosis as in rye.

Here we analyzed the drive of *Ae. speltoides* B chromosomes and provide direct insight into its cellular mechanism. We first employed comparative genomics to identify a B-specific repeat, which we used to track the B chromosomes during microgametogenesis. It was found that sister chromatid nondisjunction and an asymmetric spindle during first pollen grain mitosis are components of the B chromosome accumulation process. To determine the B accumulation rate in sperm nuclei, we developed a novel flow cytometric approach that allowed the differentiation between vegetative and sperm nuclei as well as the quantification of B chromosomes inside the nuclei. Utilizing this method, we found that independent of the number of B chromosomes present in the mother plant, Bs accumulate in the generative nuclei to more than 93%. The prerequisites for the drive process seems to be common in Poaceae, thus enabling independent origins of Bs in different lineages within the family.

## Results

### Identification of a conserved B-specific repeat suitable to trace B chromosomes in *Ae. speltoides*, *Ae. mutica* and rye

To find a B-specific marker suitable to track segregation of *Ae. speltoides* Bs during microgametogenesis by FISH, genomic 0B and +B DNA were paired-end Illumina sequenced and the corresponding reads were characterized by comparative similarity-based clustering using the *RepeatExplorer* pipeline (Novák et al., 2013). Highly and moderately abundant repeats represented by clusters containing at least 0.01% of analyzed reads were found to make up about 77% of the genome. They were mostly composed of LTR-retrotransposons, transposons and satellite repeats, with overall proportions of individual repeat types being similar in the 0B/+B genotypes (Suppl. Table 1). Comparative cluster analysis revealed four tandem repeats with monomers ranging from 86 to 1090 bp, which were found to be potentially enriched on B chromosomes (Table 1, Figure 1A-B). The B-specific candidate repeats were PCR amplified from +B genotype, cloned and verified by sequencing. In contrast to AesTR-183 and AesTR-205, AesTR-126 and AesTR-148 could also be amplified by using 0B genotype DNA as template (Suppl. Figure 1). FISH with cloned probes resulted in an exclusively B-specific localization of AesTR-183 and AesTR-205, while AesTR-126 and AesTR-148 signals were present on A and B chromosomes (Figure1C-D). In all analyzed populations, a B-specific distribution of AesTR-183 hybridization signals was found along the long B chromosome arm, while AesTR-205 clusters occur on both arms (Figure 1D). *Ae. speltoides* from the Katzir population possess in addition a morphological variant of the B, presumably lacking the terminal region of the long arm (Ruban et al., 2014). In this metacentric B variant, AesTR-183 repeats are present on both arms (Suppl. Figure 2).

**Table 1.**
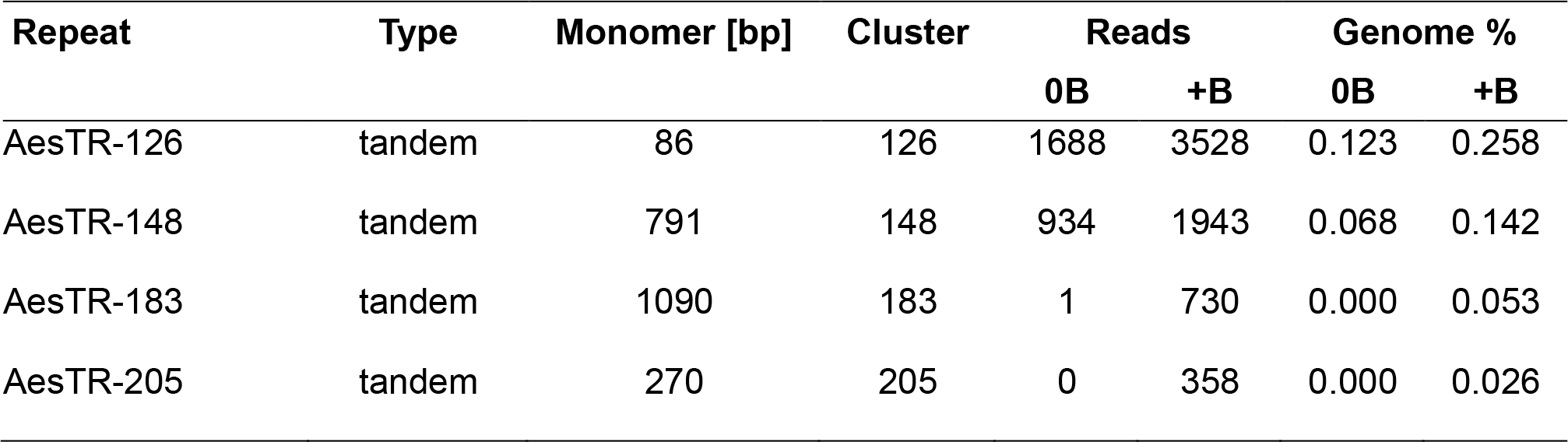
Repeats significantly enriched on the B chromosomes of *Ae. speltoides.*

| Repeat | Type | Monomer [bp] | Cluster | Reads |  | Genome % |  |
| --- | --- | --- | --- | --- | --- | --- | --- |
|  |  |  |  | 0B | +B | 0B | +B |
| AesTR-126 | tandem | 86 | 126 | 1688 | 3528 | 0.123 | 0.258 |
| AesTR-148 | tandem | 791 | 148 | 934 | 1943 | 0.068 | 0.142 |
| AesTR-183 | tandem | 1090 | 183 | 1 | 730 | 0.000 | 0.053 |
| AesTR-205 | tandem | 270 | 205 | 0 | 358 | 0.000 | 0.026 |

**Figure 1.**
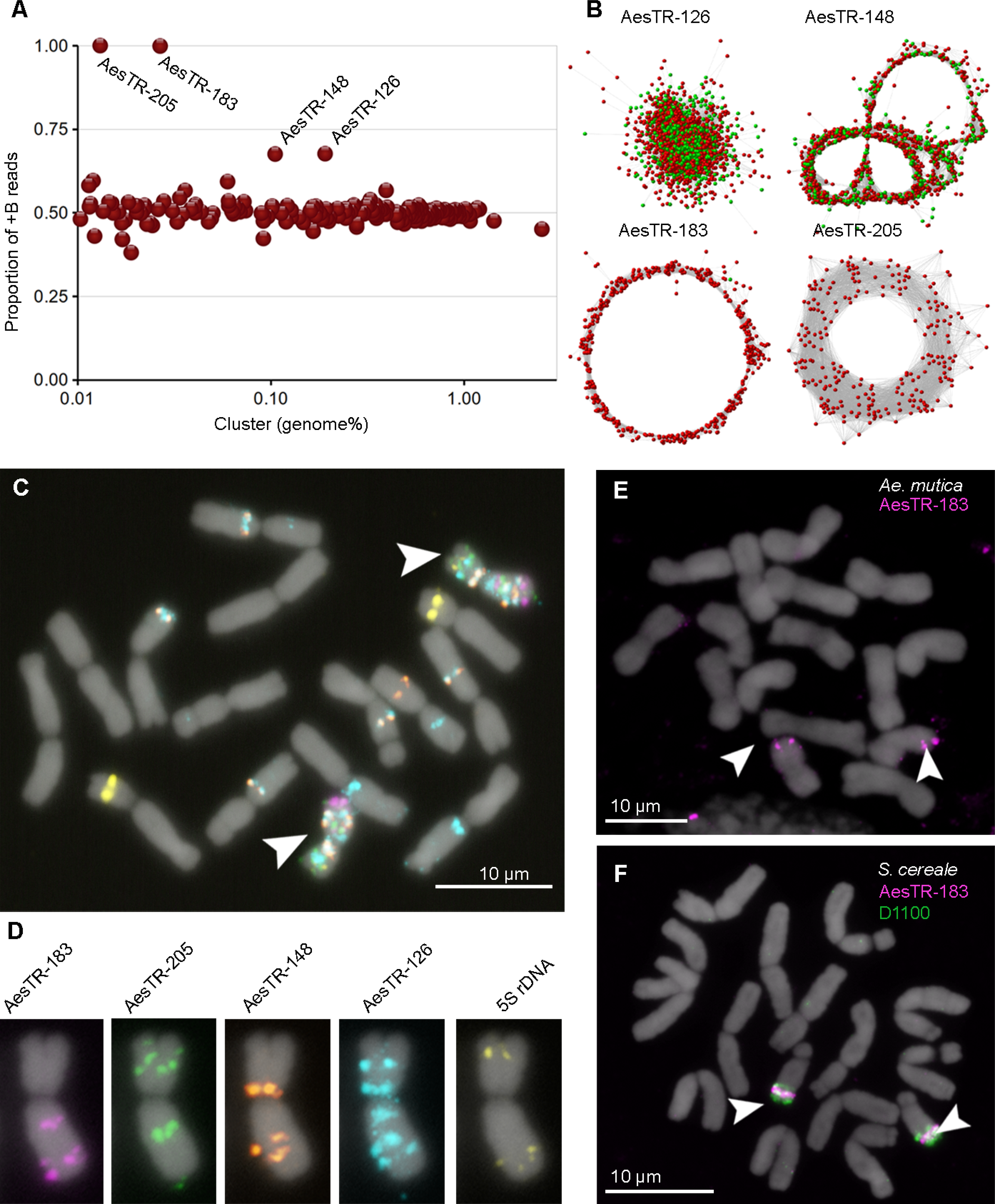
Identification and characterization of B chromosome enriched sequences. **(A)** *In silico* identification of repeat clusters containing B-enriched sequences. The enrichment is revealed by calculating the ratio of reads derived from +B genotype to the total number of reads (+B and 0B) in the cluster. Clusters showing partial enrichment (AesTR-126 and 148) or specificity to B chromosomes (AesTR-183 and 205) are labelled. **(B)** Graph representation of read similarities in B-enriched repeat clusters. Individual reads from 0B and +B plants are represented by green and red nodes, respectively. Nodes are placed according to sequence similarity, where similar sequences are close together and connected with edges (gray lines). Circular or globular structure of these graphs is typical for repeats with tandem arrangement in the genome. **(C-F)** Chromosomal distribution of B-enriched repeats in *Ae. speltoides, Ae. mutica* and *S. cereale*. **(C)** Localization of AesTR-126, AesTR-148, AesTR-183, AesTR-205 and 5S rDNA on mitotic metaphase chromosomes of *Ae. speltoides* from Tartus (Syria) revealed by FISH. **(D)** Showing the hybridization patterns of FISH probes individually. (E) Co-localization of AesTR-183 (magenta) and the *S. cereale* B-specific probe D1100 (in green) in *S. cereale* with 2Bs. **(F)** Localization of AesTR-183 in *Ae. mutica* with 2Bs. The Bs are marked with arrowheads.

Notably, the tandem repeat AesTR-183 with a monomer length of 1090 bp displayed also B-specific signals in the closely related species *Aegilops mutica* (Figure 1E). In +B rye, AesTR-183 signals are localized in the nondisjunction-controlling region of the B chromosome (Figure 1F). To test the presence of this repeat on the Bs of a more distantly related grass species, AesTR-183 probe was hybridized to a *Lolium perenne* accession carrying two Bs. However, no B-specific signal was found (data not shown). Hence, three species from the Triticeae encode a conserved B-specific repeat that can be used for the identification of Bs. BLAST (Altschul et al., 1990) search revealed 82% similarity between a part of AesTR-183 and a 489 bp long region on chromosome 1B of *Triticum aestivum* (accession LS992081.1). To estimate the relationships of corresponding sequences, a phylogenetic tree was constructed based on sequences of PCR amplified AesTR-183 monomers from *Ae. speltoides*, *Ae. mutica* and rye, including the similar sequence from wheat chromosome 1B (Suppl. Figure 3). The obtained phylogenetic tree placed the analyzed sequences in three clades according to the species of origin. The much longer branch lengths in the case of rye-derived sequences indicates higher substitution rate. Surprisingly, the sequence from wheat chromosome 1B appeared as sister group to the rye B-located sequences, while it would rather be expected to cluster together with *Ae. speltoides*-derived sequences due to closer evolutionary relationships of wheat and *Ae. speltoides* genomes (Bernhardt et al., 2017). Interestingly, the diversity of AesTR-183 repeat monomers in *Ae. mutica* was much lower than in *Ae. speltoides* and rye.

### Flow cytometry enables to determine the accumulation of B chromosomes in sperm nuclei

To quantify the accumulation of *Ae. speltodies* B chromosomes during pollen development we applied a combination of flow cytometry, immunostaining and FISH. Differences in size and shape of nuclei isolated from mature pollen grains (Figure 2A) of a 0B plant allowed the separation of vegetative and sperm nuclei by plotting the fluorescence intensity (DNA content) against the forward scatter signal (FSC) representing particle size. The round-shaped and larger vegetative nuclei appear in the upper fraction while the spindle-shaped smaller sperm nuclei were found in the lower fraction (Figure 2B). To confirm the flow-cytometric separation of both types of nuclei we sorted both fractions and used the nuclei for immunolabelling with an antibody against histone H3K27me3 which was shown to be preferentially enriched in vegetative nuclei in rye (Houben et al., 2011). Indeed, an intense immunolabelling of nuclei was only observed on samples derived from the upper fraction representing the vegetative nuclei (Figure 2C) but not on sperm nuclei (Figure 2D). Thus, both types of nuclei can easily be differentiated flow cytometrically.

**Figure 2.**
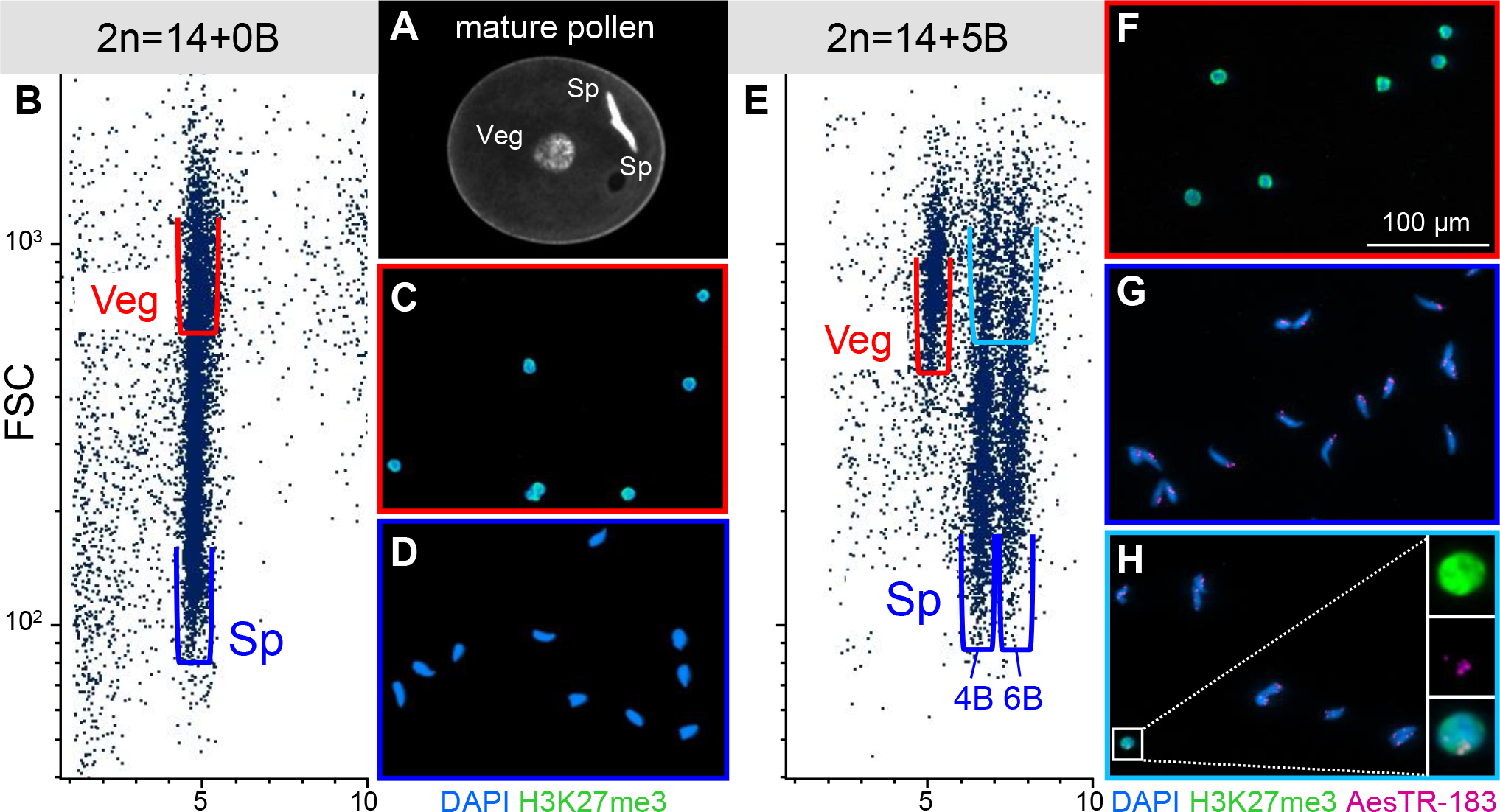
Flow cytometry showed high efficiency of B chromosome drive. **(A-D)** Vegetative (Veg, red) and sperm (Sp, blue) nuclei of 0B mature pollen were separated flow cytometrically and the purity of both fractions was verified by H3K27me3 immunostaining. **(E-F)** Vegetative (red) and sperm (blue) nuclei of +5B mature pollen were sorted and subjected to H3K27me3 immunostaining and B-specific FISH using AesTR-183. **(A)** PI stained mature pollen grain from 0B plant contained round vegetative nucleus and two spindle shaped sperm nuclei. **(B)** Dot plot displaying the relative fluorescence (DNA content; x-axis) versus forward scatter signal (particle size; y-axis) intensities. **(C)** Sorted vegetative nuclei fraction (red) was abundantly labelled by anti-H3K27me3 (in green). **(D)** Sperm nuclei fraction (blue) showed no anti-H3K27me3 signal. **(E)** Dot plot of +5B plant pollen with fraction of vegetative (red) and two fractions of different fluorescence intensities of sperm nuclei (blue). **(F)** Sorted vegetative nuclei labelled by H3K27me3 showed no B-specific FISH signals. **(G-H)** Sorted sperm nuclei after immunostainig with H3K27me3 and FISH with the B-specific probe AesTR-183. While in the lower part (blue) of the sperm nuclei fractions exclusively B-carrying sperm nuclei **(G)** were detected the upper fraction (light blue) contained also a low percentage of vegetative nuclei with B chromosomes **(H)**.

In nuclei suspensions of pollen derived from plants with Bs, additional nuclei populations were observed in the dot plots (Figure 2E, Suppl. Figure 4B-G). Depending on the number of Bs in the mother plant we detected one to three additional nuclei populations at similar forward scatter signal intensities as detected for the sperm nuclei of the 0B plants (Suppl. Figure 4A-H) but with a higher fluorescence intensity indicating an increased DNA content due to the presence of additional B chromosomes. The number of Bs represented by the individual populations could be estimated by the distance between the fractions of vegetative nuclei, which appeared unchanged, and the corresponding fraction of sperm nuclei. In case of a 1B plant, one additional population of sperm nuclei appeared at a position that indicates the presence of two B chromosomes. However, the majority of sperms showed no Bs (Suppl. Figure 4B). This observation is explainable by the fact that the B chromosome appears as a univalent during metaphase I leading either to laggards mostly being excluded from the new daughter nuclei or to the migration of the single B chromosome to one of the two poles. In both cases the second meiotic division occurs normally, only in the latter case two of the four spores contain one B chromosome each (Mendelson and Zohary, 1972) (Suppl. Figure 5). Consequently, sperm nuclei formed after two rounds of mitotic divisions contain no or two B chromosomes. In our experiments we detected 25% of sperm nuclei containing 2Bs (Table 2). In pollen of 2B plants almost exclusively sperm nuclei with two Bs were detected (Suppl. Figure 4C). In this case the two Bs frequently form bivalents during meiosis I and segregate normally. At the end of meiosis spores with one B chromosome each are formed (Mendelson and Zohary, 1972) which result in sperm nuclei with two chromosomes (Suppl. Figure 5). 3B plants produced pollen with more or less equal sized fractions of sperm nuclei with two and four B chromosomes (Suppl. Figure 4D) similarly as plants with 5Bs where nuclei with four and six B chromosomes (Suppl. Figure 4E) were detectable. For 3B plants the occurrence of univalents, bivalents and trivalents was described (Mendelson and Zohary, 1972). Resolving of these pairing structures preferentially results in spores with either one or two B chromosomes finally leading to sperm nuclei with two or four Bs (Suppl. Figure 5). However, with a much lower frequency also sperm nuclei with no and six Bs could be detected (Suppl. Figure 4D). In plants with 5 Bs the number of potential pairing configurations further increases. However, preferably either two or three Bs migrate to opposite poles during meiosis I finally resulting in sperm nuclei with either four or six Bs. In pollen of plants with 4Bs and 6Bs preferentially sperm nuclei with the same number of B chromosomes as in the corresponding mother plant were produced and with a lower frequency nuclei with two and six or four and eight Bs, respectively (Suppl. Figure 4F-G). Plants with four or six Bs preferentially perform regular segregations during meiosis independent on the pairing configuration (bivalents up to hexavalents) (Mendelson and Zohary, 1972). Deviations from this normal segregation may lead to a decrease or increase of B chromosomes in sperm nuclei compared to the situation in the mother plant (Table 2).

**Table 2.** Quantification of B chromosome accumulation in sperm nuclei.

| Mother plant | Drive (%)<sup>1</sup><br>(number of nuclei<br>counted) | Frequency of Bs in sperm nuclei (%)<sup>2</sup> |  |  |  |  |
| --- | --- | --- | --- | --- | --- | --- |
|  |  | 0B | 2B | 4B | 6B | 8B |
| 1B | 94.4 (407) | 75.1 | 24.9 |  |  |  |
| 2B | 93.9 (770) | 1.9 | 98.1 |  |  |  |
| 3B | 98.3 (754) | 6.5 | 48.0 | 39.5 | 5.9 |  |
| 4B | 97.9 (660) | 2.1 | 14.4 | 74.2 | 9.2 |  |
| 5B | 96.3 (589) | 1.4 |  | 57.3 | 41.4 |  |
| 6B | n.d. | 2.5 |  | 24.6 | 61.7 | 11.2 |

<sup>1</sup> Bs drive efficiency calculated based on vegetative nuclei without Bs after flow sorting, immunostaining and FISH
<sup>2</sup> frequencies were quantified flow cytometrically

To measure the efficiency of the B accumulation, vegetative and sperm nuclei from mature pollen of individual +B plants were separately sorted onto microscopic slides first (red and blue frames in Figure 2E). Application of immunolabelling using anti-histone H3K27me3 to identify the vegetative nuclei in combination with FISH using the B-specific repeat AesTR-183 to score for the presence of Bs (Figure 2F-G) confirmed the purity of the individual populations. In order to evaluate if the B chromosome-containing fractions with a FSC signal intensity similar to that of the vegetative nuclei (light blue frame in Figure 2E and Suppl. Figure 4B-H) exclusively contain sperm nuclei or also vegetative nuclei with Bs we sorted the upper (light blue) and lower (dark blue) fractions of sperm nuclei separately. Only the upper sperm nuclei fractions (light blue) contained additionally a low percentage of vegetative nuclei harboring Bs (Figure 2H) in a range of 1.7% to 6.1%. Based on these cytological evaluations in combination with the flow cytometric profiles we were able to precisely determine the transmission of Bs to sperm nuclei (Table 2). Independent of the number of Bs present in the mother plant Bs accumulate in the sperm nuclei with more than 93% as a result of an unequal segregation of Bs during the first pollen mitosis.

Since in rye a structural change of the B chromosome impaired its ability to drive (Endo et al., 2008; Müntzing, 1945), we included *Ae. speltoides* plants from the Katzir population with a metacentric B variant in our analysis (Suppl. Figure 2B). However, based on flow cytometry, this B-variant shows a comparable frequency of accumulation in sperm nuclei (Suppl. Figure 4H). Hence, the changes involved in the formation of this B-variant did not affect the process behind the control of its preferential transmission to sperm nuclei.

### B chromosomes preferentially distribute in the generative nucleus of the bicellular pollen grain

To obtain visual insights into the cellular mechanism of B chromosome drive in *Ae. speltoides*, we analyzed the dynamics of chromosomes at microgametogenesis. Anther tissue sections were used to maintain the natural arrangement of chromosomes and to avoid pollen wall background signals. The B-specific probe AesTR-183 was used to trace the segregation of Bs during pollen formation.

Products of the pollen mitosis I were analyzed first. Due to differences in the chromatin condensation and size of generative and vegetative nuclei (Borg et al., 2009; Tanaka, 1997), both types of nuclei are easily distinguishable. Condensed generative nuclei locate closely toward the pollen wall. 120 bicellular pollen grains of plants possessing 2Bs were analyzed. In 85% of pollen, the Bs were observed in the generative nucleus only (Figure 3A). Micronuclei containing Bs were found in close vicinity to the generative nucleus in 3.3% of pollen (Figure 3B). 3.3% of pollen revealed B-specific signals in vegetative nuclei only (Figure 3C). 1.7% of pollen revealed Bs in the generative nucleus and in micronuclei (Figure 3D). 6.7% of pollen showed B-specific signals in both, generative and vegetative nuclei (Figure 3E). In all 10 analyzed mature tricellular pollen grains, B-specific signals were never observed in vegetative nuclei, while all of the sperm nuclei revealed signals; indicating a normal segregation of Bs at second pollen grain mitosis (Figure 3F). Thus, Bs accumulate in the generative nucleus during the first pollen mitosis. The observed values of the B chromosome accumulation in tissue sections are in close agreement with the flow cytometric data.

**Figure 3.**
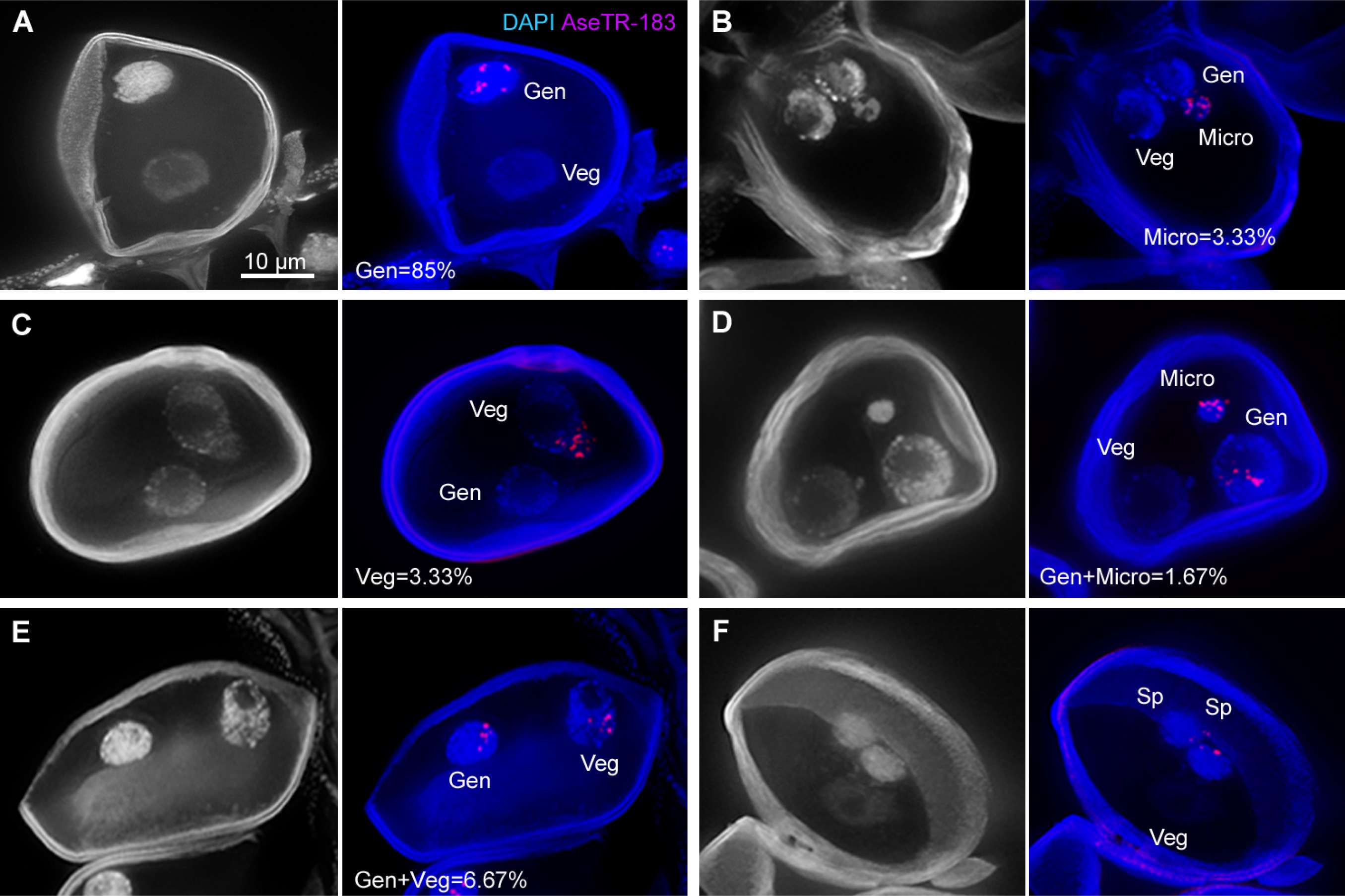
B chromosomes of *Ae. speltoides* preferentially accumulate in the generative nucleus during first pollen grain mitosis. Tissue sections after FISH with the B-specific repeat AesTR-183 (magenta). **(A)** 85% of binucleated pollen possess Bs in generative nuclei (Gen) only. **(B)** 3.33% of binucleated pollen possess Bs in micronuclei (Micro) only. **(C)** 3.33% of binucleated pollen possess Bs in vegetative nuclei (Veg). **(D)** 1.67% of binucleated pollen possess Bs in generative and micronuclei. **(E)** 6.67% of binucleated pollen possess Bs in generative and vegetative nuclei. **(H)** In mature pollen both sperm nuclei possess Bs. DAPI is showen in blue.

### Directed nondisjunction of *Ae. speltoides* Bs is accompanied by unequal cell division during pollen mitosis

To decipher the mechanism behind the preferential accumulation of B chromosomes in the generative nucleus, tubulin and CENH3 antibodies were applied at all stages of the first pollen mitosis of B chromosome-containing plants. The centromere-specific histone variant H3 (CENH3) was selected because it was shown that its loss results in the failure of chromosome segregation in mammals (Howman et al., 2000) and plants (Sanei et al., 2011). The centromeres of all anaphase A and B chromosomes were equally CENH3 labeled and revealed an interaction with microtubules. During early anaphase, the peripheral generative spindle bundle is blunt and the pole is nearly in contact with the pollen cortex. By contrast, the interior spindle bundle is long and pointed (Figure 4A). In late anaphase, a nondisjoined lagging chromosome with attached spindle fibers was observed (Figure 4B). During telophase, the generative nucleus contained a higher number of CENH3 signals indicating the presence of Bs (Figure 4C). Comparing the distance between the spindle midzone and the centromere positions of the generative (6.82 μm) versus the vegetative nucleus (7.97 μm) revealed that in most cases the spindle midzone is more proximal to the generative nucleus (n = 10, t-test P<0.05) (Figure 4D).

**Figure 4.**
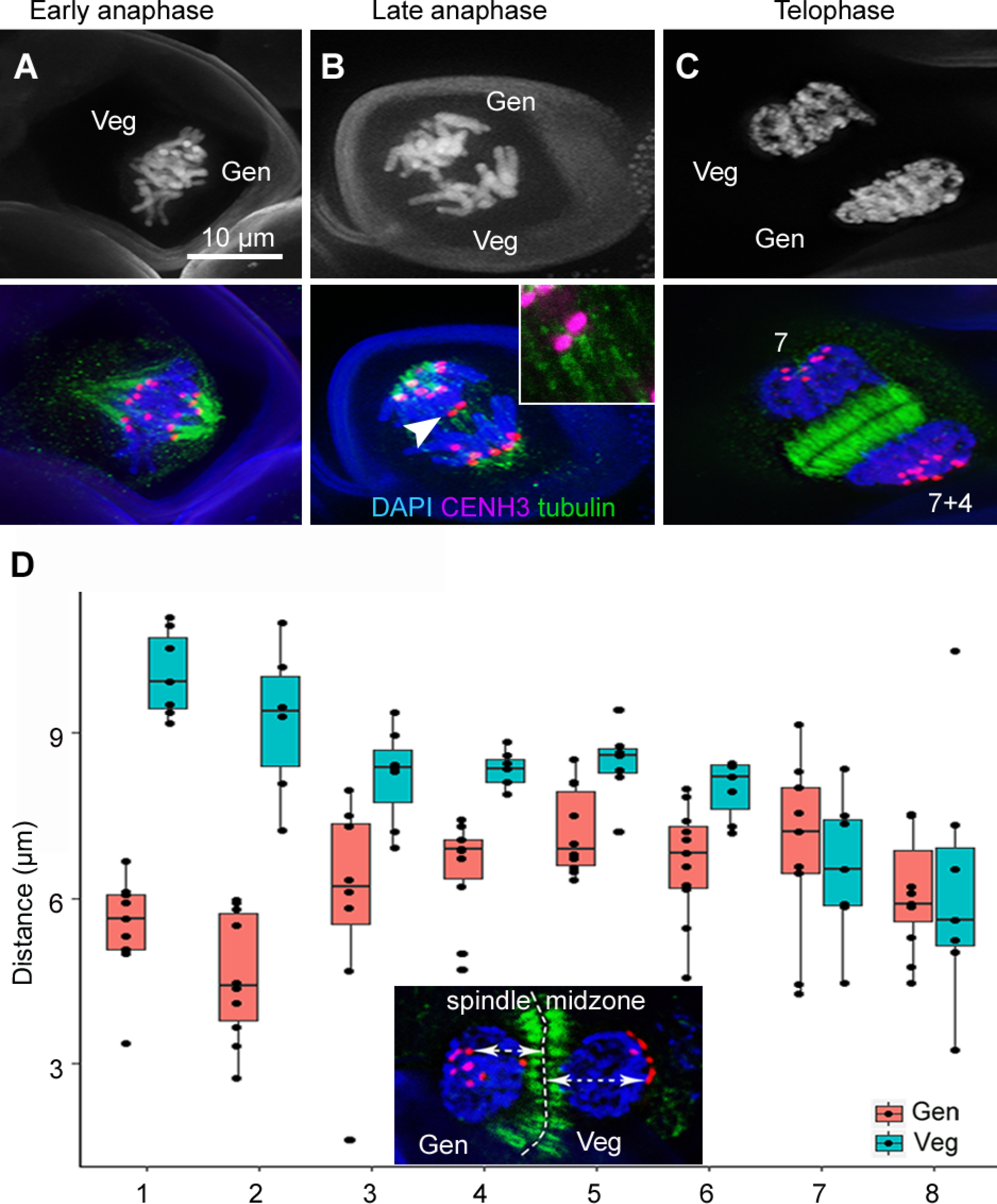
The accumulation of Bs during pollen mitosis of *Ae. speltoides* +2B. **(A-C)** CENH3 (magenta) and α-tubulin (green) immunolabelling patterns at different stages of first pollen mitosis. **(A)** Formation of asymmetric spindle at early anaphase. The generative spindle (Gen) is short and blunt and the vegetative spindle (Veg) is long and sharp. **(B)** At late anaphase, nondisjunction of lagging chromosomes with positive CENH3 signals attached to the tubulin (insert shows further enlarged CENH3 signals). **(C)** Pollen at late anaphase forming an asymmetric spindle midzone. Note the unequal number of CENH3 signals. **(D)** The dots indicate the distance between the spindle midzone and the individual centromeres of the generative nucleus (red) or vegetative nucleus (blue). X-axis: individual cells, Y-axis: distance.

In order to quantify the centromere size as a means for centromere activity (Zhang and Dawe, 2012), the size of immunosignals was determined after anti-CENH3 labeling and B-specific *in situ* hybridizations of somatic mitotic +B metaphase cells. No major differences in CENH3 signal size were found when we compared A and B centromeres, although the amount of CENH3 slightly varied between the individual chromosomes (Figure 5A-B). Similarly, we did not observe differences in CENH3 signal size using meiotic prophase cells. Thus, a different centromere activity of B chromosomes could be excluded as the reason for the nondisjunction of Bs, since the centromeres of all chromosomes are CENH3-positive and interact with tubulin.

**Figure 5.**
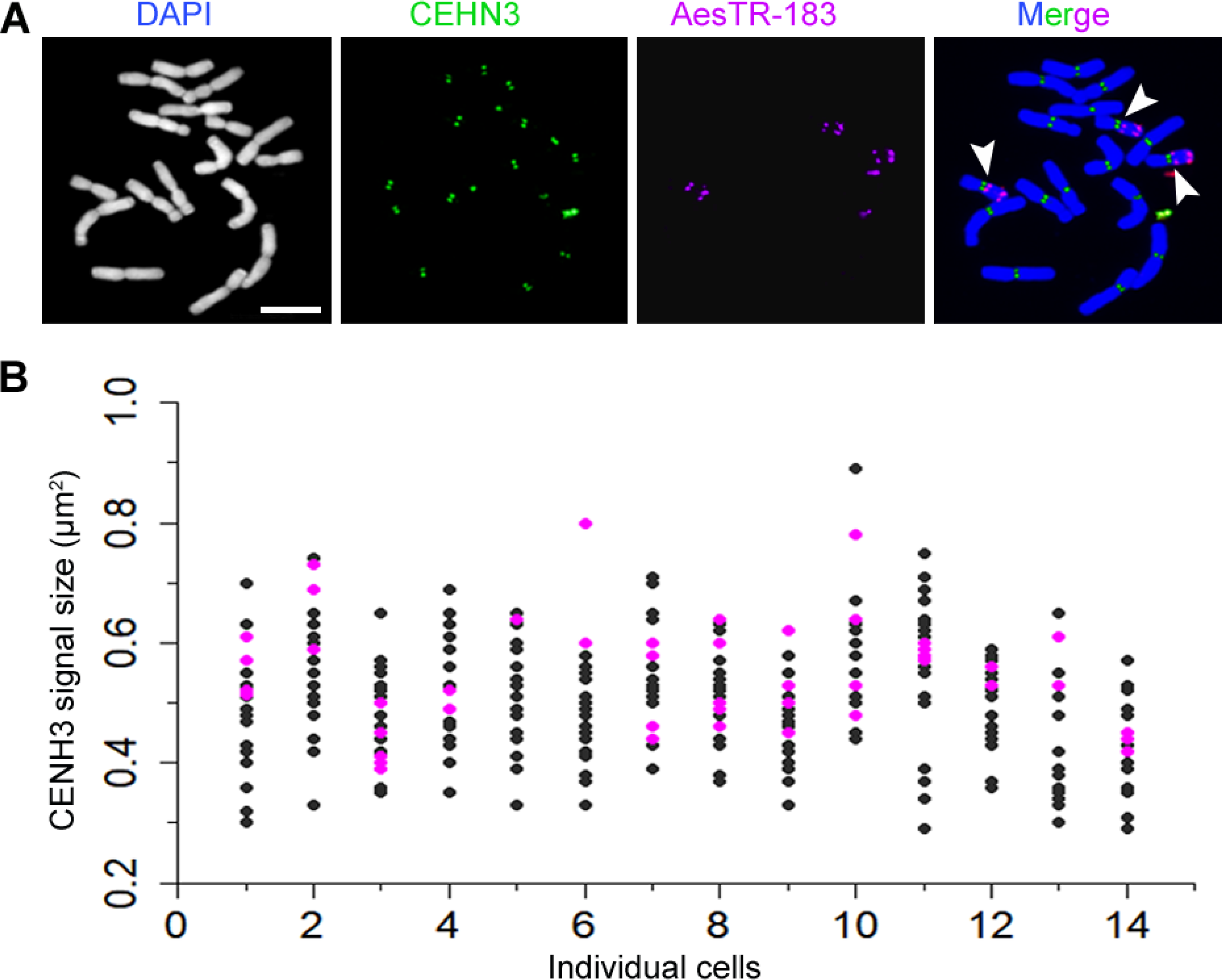
Size of CENH3 immunosignals does not differ between mitotic A and B chromosomes of *Ae. speltoides*. **(A)** Mitotic metaphase of a 3B *Ae. speltoides* plant after CENH3 immunostaining (green) and FISH with the B-specific probe AesTR-183 (magenta). Bs are marked with arrowheads. Scale bar equals 10 μm. **(B)** Each dot represents the CENH3 signal size of an individual sister chromatid (As in black, Bs in magenta). X-axis: individual cells, Y-axis: CENH3 signal size.

### The (peri)centromere composition differs between A and B chromosomes of *Ae. speltoides*

Next, we asked whether the sequence composition differs between A and B (peri)centromeres. Eleven (peri)centromere-specific repeats, originally described for wheat were identified in *Ae. speltoides.* The corresponding FISH probes were generated by PCR using genomic DNA of *Ae. speltoides* carrying B chromosomes (Suppl. Table 2,) and hybridized to mitotic metaphase chromosomes of +B plants (Figure 6, Suppl. Figure 6, Table 3). Four of the eleven probes (RCS2, 6C6-3, 6C6-4 and CRW2) resulted in signals of similar intensities on A and B chromosomes (Figure 6A, Suppl. Figure 6A-C). Four probes (Quinta-LTR, 192 bp, Weg 1-LTR and CCS1) showed stronger hybridization signals on the A chromosomes (Figure 6B, Suppl. Figure 6D-F), and pBs301 and the centromeric satellite repeat (CenSR) co-localized with the pericentromeres of As but were not detectable on Bs (Figure 6C, Suppl Figure 6G). In contrast, TailI revealed an enriched signal accumulation at centromeres of Bs (Figure 6D). No obvious differences in the distribution and intensity of the used (peri)centromeric probes were found between the different A chromosomes of *Ae. speltoides.*

**Table 3.** Characterization of the employed (peri)centromeric probes.

| (peri)centromeric repeat | Signal intensity and size<sup>1</sup> |  | CENH3 interaction<sup>2</sup> |  | Sequence type |
| --- | --- | --- | --- | --- | --- |
|  | As | Bs | ChIP<br>(original<br>paper) | FISH in<br>pachytene of<br>*Ae.speltoides* |  |
| 192 bp (Ito et al. 2004) | ** | * | + | + | Ty3/gypsy LTR |
| 6C6-3 (Zhang et al. 2004) | *** | *** | + | + | Ty3/gypsy cereba gag |
| 6C6-4 (Zhang et al. 2004) | *** | *** | + | + | Ty3/gypsy cereba gag |
| CCS1 (Aragon-Alcaide et al. 1996) | *** | * | + | + | Ty3/gypsy LTRs |
| <u>Centromeric satellite repeats</u><br>(CenSR) (Li et al., 2013) | ** | - | - | - | centromeric satellite |
| CRW2 (Liu et al. 2008) | *** | *** | + | + | Ty3/gypsy cereba LTR |
| pBs301 (Cheng & Murata 2003) | ** | - | + | - | Ty3/gypsy family |
| RCS2 (Liu et al. 2008) | *** | *** | + | - | Ty3/gypsy family |
| Quinta-LTR (Li et al., 2013) | ** | * | + | + | Ty3/gypsy Quinta LTR |
| Tail-family (Kishii et al., 2001) | * | *** | n.d. | + | tandem repeat |
| Weg 1-LTR (Li et al., 2013) | ** | * | + | + | Ty3/gypsy Weg 1-LTR |

1 “\*” represents the appearance of signal, “-” means no signal detected
2 “+” represents (peri)centromere colocalized with CENH3 antibody, “-” means no interaction, “n.d.” no data available

**Figure 6.**
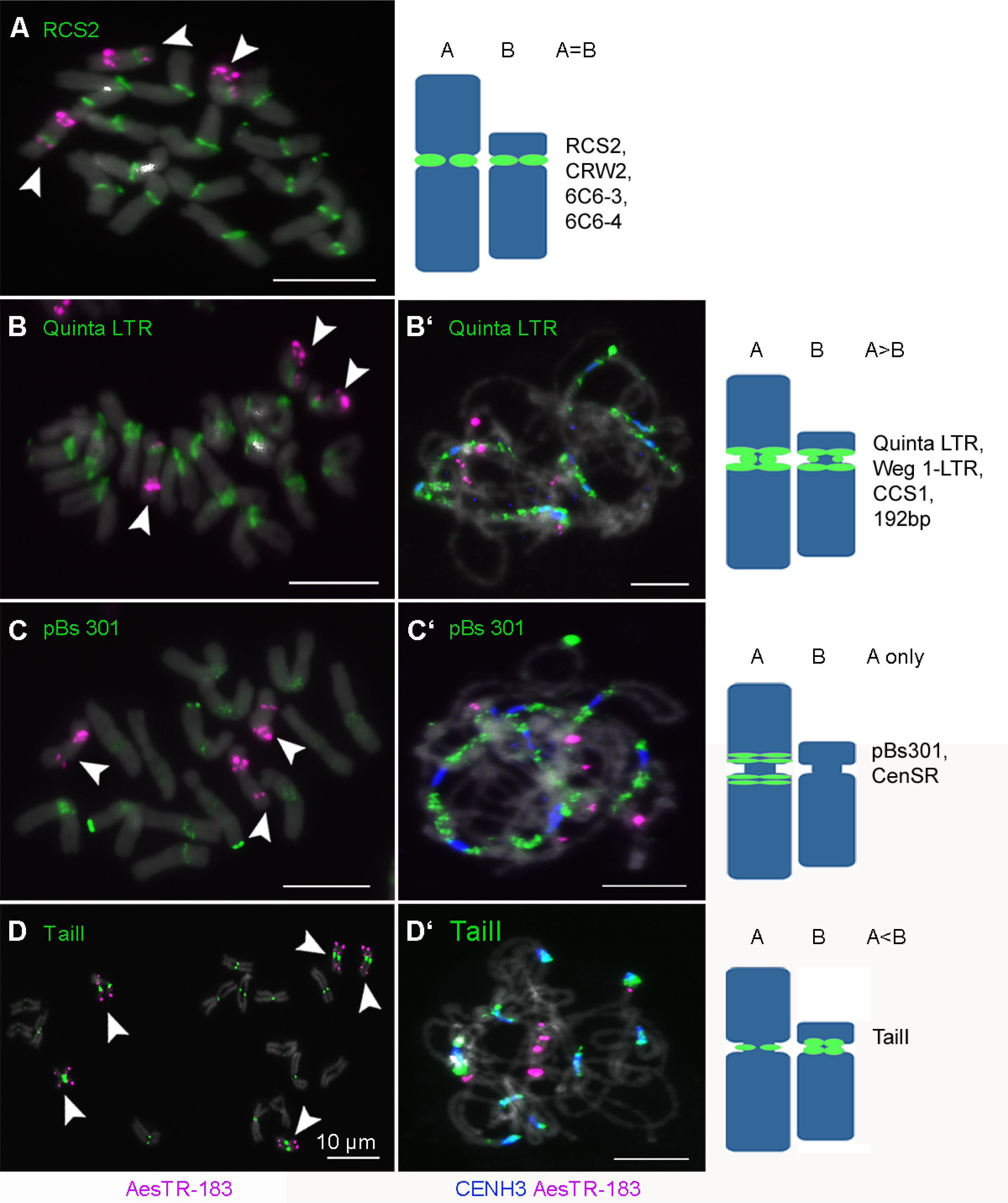
Sequence composition of *Ae. speltoides* A and B (peri)centromeres. **(A-D)** FISH patterns of RCS2, Quinta-LTR, pBs301 and TailI (green) as well as of the B-specific repeat AesTR-183 (magenta) on mitotic metaphase chromosomes. White arrows indicate Bs. **(B’-D’)** Meiotic pachytene chromosomes labelled with the corresponding (peri)centromeric probes (except for RCS2) and with antibodies against CENH3 (blue). Schematic chromosomes represent typical (peri)centromeric distributions of the corresponding sequences on A and B chromosomes. Hybridization patterns of the remaining (peri)centromeric sequences are shown in Suppl. Figure 6. DAPI staining shown in grey. Scale bars equal 10 μm.

To evaluate the centromeric localization of the identified repeats in more detail we used naturally extended pachytene chromosomes for anti-CENH3 staining and subsequent FISH with the corresponding repeats. AesTR-183 was additionally used to distinguish between A and B chromosomes. Quinta-LTR, 192 bp, Weg 1-LTR (Figure 6B’, Suppl. Figure 6D’-E’) signals partly overlapped with CENH3-marked chromatin and extended into pericentromeric regions. CCS1 completely overlapped with CENH3 signals in A chromosomes showed reduced centromere labeling in B chromosomes (Suppl. Figure 6F’). No overlapping with CENH3-signals was found for the A chromosome-specific pBs301 and CenSR (Figure 6C’, Suppl. Figure 6G’). TailI coincided with CENH3 and showed stronger centromere signals on Bs than on As (Figure 6D’). Although nine of the eleven (peri)centromeric candidate sequences co-localize with CENH3 signals, CENH3 never encompassed fully the area covered by centromere repeats. We conclude that all A centromere sequences except pBs301 and the CenSR are also present in the centromeres of the Bs. Hence, an intraspecific origin of the B centromere is likely. However, the contrasting abundance of Quinta-LTR, 192 bp, Weg 1-LTR and CCS1 signals indicates that the (peri)centromere of the B diversified.

## Discussion

### AesTR-183, a conserved B chromosome-specific repeat

The high copy-repeat fraction of the *Ae. speltoides* genome is mostly composed of LTR-retrotransposons, transposons and satellite repeats, with overall proportions of individual repeat types being similar in the 0B/+B genotypes. Among four identified B-located repeats in *Ae. speltoides* only AesTR-183 and AesTR-126 showed similarity to already known repeats in other species according to BLAST results. AesTR-126 appeared to be identical to a part of the *T. aestivum* clone pTa-713, a repetitive sequence positively tested in FISH on wheat chromosomes (KC290900.1) (Badaeva et al., 2015). AesTR-183 shows 84% similarity to the rye B-specific tandem repeat Sc26c38 which occurs within the nondisjunction control region of the rye B (Klemme et al., 2013). In addition, we discovered an AesTR-183-like repeat in the Bs of *Ae. mutica.* Unknown is, whether the Bs of *Ae. speltoides* and *Ae. mutica* possess like rye a repeat-enriched nondisjunction control region. The existence of a B chromosome conserved repeat is intriguing and provokes the question whether the AesTR-183 repeat has acquired a function in Bs. Possibly, the ancestor of all three species possessed an AesTR-183-positive B chromosome, which evolved further in rye and *Aegilops*, but kept the AesTR-183 sequence. The phylogenetic analysis of AesTR-183 repeat monomer sequences rather support this theory, as the divergence of B-specific repeat in this case coincides with divergence of the species themselves (Bernhardt et al., 2017). The similar but much shorter sequence detected on wheat chromosome 1B does not follow this pattern and seems to be independent from the B-specific sequences.

### A combination of nondisjunction and unequal spindle formation at first pollen mitosis might promote the accumulation of Bs in sperm nuclei

We provide direct insight into the cellular basis of the drive mechanism of the B chromosome. The flow-cytometrically estimated frequencies of B chromosome accumulation correspond with the cytologically determined one using tissue sections of developing pollen grains (93.9% versus 95%). However, one advantage of the cytological approach is the observation of micronuclei, which escaped from detection in the flow cytometrical approach. On the other hand, flow cytometry allows the rapid analysis of a large number of pollen nuclei of different plants. This approach seems to be applicable also in other plant species with B chromosomes provided the structural and morphological differences between vegetative and sperm nuclei allow a clear differentiation, and the size of the B chromosomes is large enough to separate individual fractions with different B chromosome numbers.

More than 93% of sperm nuclei accumulated Bs independent on the number of Bs present in the mother plant. In +6B plants, sperm nuclei with up to 8Bs were found. Eight reflects the maximum number of Bs detected in *Ae. speltoides*. Thus, embryos with a B number above 8 are not formed or do not develop due to the negative effect of Bs in a high number. It would be of interest to investigate pollen of +8B plants to see if at least sperm nuclei with more than eight Bs are produced. However, plants with 8Bs were not found during this study and are probably male sterile.

An important hint regarding the mechanism of chromosome accumulation comes from our finding that the microtubule spindle of *Ae. speltoides* is asymmetrical during first pollen mitosis, in accordance with previous studies in other species (Banaei-Moghaddam et al., 2012; Borg et al., 2009; Jin et al., 2005; Tanaka, 1997; Twell, 2011). Considering the asymmetric geometry of the spindle at first pollen grain mitosis, it is likely, that the inclusion of Bs into the generative nucleus is caused by the fact that the equatorial plate is nearer to the generative pole as suggested by Jones (1991). Spindle asymmetry as a component of the B chromosome drive process has been suggested also for the lily *Lilium callosum* (Kimura and Kayano, 1961), the Asteraceae *Crepis capillaris* (Rutishauser and Rothlisberger, 1966) and the grasshopper *Myrmeleotettix maculatus* (Hewitt, 1976). Alternatively, due to a higher pulling force on the B centromere toward the generative pole, Bs may preferentially accumulate in the generative nucleus (Banaei-Moghaddam et al., 2012).

Mitotic nondisjunction and lagging chromatids could be explained by centromere inactivity (Sanei et al., 2011). However, as the centromeres of lagging Bs show no reduction of CENH3 signals and interact with spindle microtubules, centromere inactivity is unlikely the reason behind the nondisjunction process. In rye, the failure to separate CENH3-positive B sister pericentromeres during pollen grain mitosis is under the control of the nondisjunction control region (Banaei-Moghaddam et al., 2012; Endo et al., 2008). Also in maize, centromere function and B chromosome nondisjunction are two independent processes, but the region required for nondisjunction resides in the centromeric region (Han et al., 2007). It is likely, that in *Ae. speltoides* and rye the cohesion between B sister chromatids during first pollen mitosis is stronger than the microtubule traction force required for separation of chromatids. At second pollen grain mitosis, B sister chromatids disjoin and a normal segregation of Bs occurs. The proposed mechanism of B chromosome accumulation is summarized in Figure 7.

**Figure 7.**
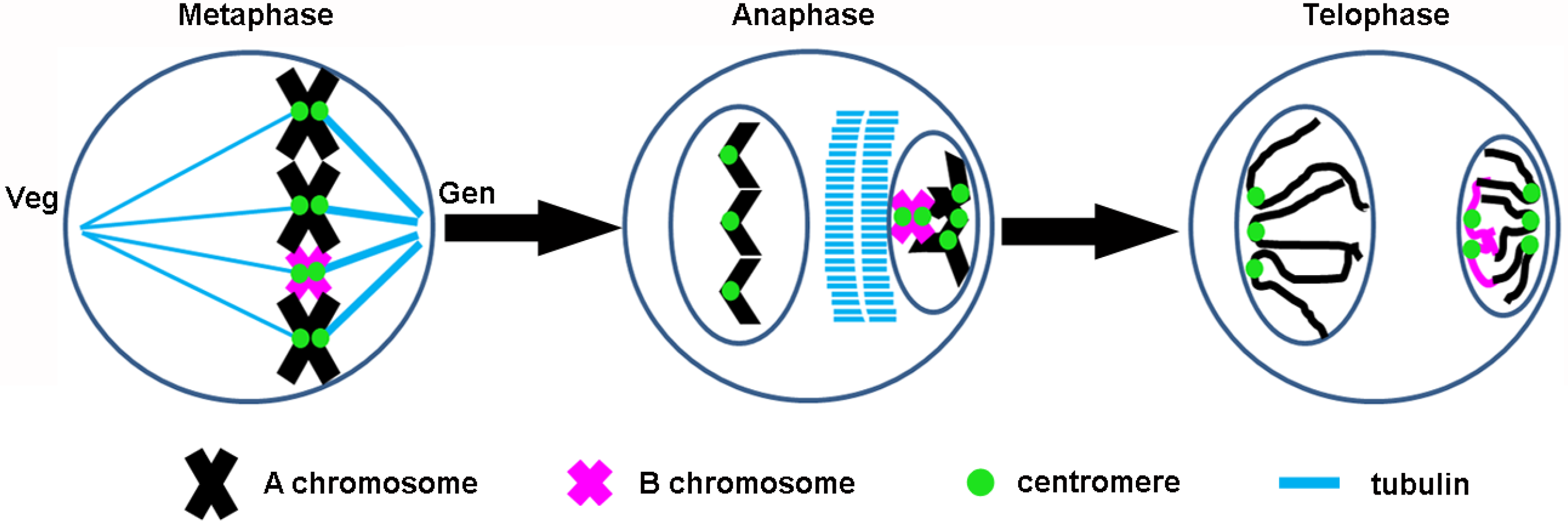
Model to explain the accumulation of *Ae. speltoides* B chromosomes during male gametophyte development. At first pollen mitosis, metaphase chromosomes are located toward the pollen cortex and an asymmetric spindle is formed. The peripheral generative (Gen) spindle bundle is blunt and the interior spindle bundle toward the vegetative pole (Veg) is longer and pointed. In contrast to the separated A sister chromatids, the cohesion of B chromatids stays unresolved and Bs migrate preferentially to generative nuclei. The placement of Bs toward the generative nucleus is probably realized because the tubulin midzone is closer to the generative pole and lagging Bs are included in the generative nucleus as the nuclear membrane is formed. Alternatively, a higher spindle tension force toward the generative pole might preferentially pull the Bs to the generative nucleus. Unlike As, nondisjoined B chromatids locate preferentially at a position close to the spindle midzone.

Notably, the B chromosome accumulation mechanism in *Ae. speltoides* and rye is similar. In both species, nondisjunction of CENH3-positive B sister chromatids occurs and an asymmetric spindle during first pollen grain mitosis are likely components of the accumulation process. Assuming that the Bs of rye and *Ae. speltoides* originated independently, the comparable drive mechanisms in both species likely evolved in parallel.

### Similarities in the (peri)centromere composition of As and Bs suggest an intraspecific origin of B chromosomes in *Ae. speltoides*

The observed generally similar sequence composition of the A and B centromeres of *Ae. speltoides* could be considered as evidence of an intraspecific A chromosome-derived origin of the B chromosome. Nine of eleven tested (peri)centromeric sequences were shared by both types of chromosomes. Only CenSR and pBs301 did not reveal a hybridization signal in the B centromeres. However, copy number differences assumed by different FISH signal sizes were also found for the repeats Quinta-LTR, 192 bp, Weg 1-LTR, CCS1 and Taill. One might imagine that the B centromere evolved from a standard A chromosome centromere and the repeats rearranged in the newly formed B chromosome. A comparable situation was also observed for the B chromosome centromeres of the daisy *Brachycome dichromosomatica* (Houben et al., 2001), maize (Jin et al., 2005), rye (Banaei-Moghaddam et al., 2012) and the grasshopper *Xyleus discoideus angulatus* (Bernardino et al., 2017). Likely is that the B-specific centromere composition is functionally involved in the regulation of chromosome segregation to ensure the maintenance of Bs in natural populations.

### Conclusions

Drive is the key for understanding supernumerary B chromosomes. Nondisjunction of CENH3-positive, tubulin interacting B sister chromatids and an asymmetric spindle during first pollen grain mitosis are likely components of the B chromosome drive process. A quantitative flow cytometric approach demonstrated that independent of the number of Bs present in the mother plant, B chromosomes accumulate in the generative nuclei with more than 93%. The prerequisites for the drive process seems to be common in Poaceae, thus enabling independent origins of Bs in different lineages within the family.

## Methods

### Plant materials and cultivation

Following accessions were used: *Aegilops speltoides* from Syria (PI 487238, USDA-ARS, USA) and Israel (Katzir, TS 89; Technion, 2-36; Ramat Hanadiv, 2-46, Institute of Evolution, University of Haifa, Israel) (Belyayev and Raskina, 2013); *Aegilops mutica* (KU-12008, Plant Germplasm Institute, Kyoto University, Japan) (Ohta and Saruhashi, 1999); *Triticum aestivum* ‘Chinese Spring’ (TRI 12922, IPK-Gatersleben Genebank, Germany); *Secale cereale* inbred line 7415 (Jimenez et al., 1994) and *Lolium perenne* accession G7 with Bs (Taylor and Evans, 1977). Plants were cultivated under greenhouse conditions (16 h light, 22°C day/16°C night).

### Isolation and sequencing of genomic DNA

Genomic DNA from leaf tissue was purified with DNeasy Plant Mini Kit (Qiagen, Germany). Using the Illumina HiSeq 2500 system, we sequenced 2× 101 bp genomic DNA from 0B and 2B plants from the *Ae. speltoides* Katzir accession. In total 171 million Illumina 101 bp paired end reads were generated having an output of residues of 17 Gbp. All sequences are publicly available under PRJEB29862 (ENA).

### Identification of B-specific repeats using similarity-based read clustering

Identification of *Ae. speltoides* B chromosome-specific repeats was performed by similarity-based clustering of Illumina reads as implemented in the *RepeatExplorer* pipeline (Novák et al., 2010; Novák et al., 2013). The analysis was done in a comparative mode, using as its input a randomly sampled fraction of 1.4 million paired-end reads from 0B and +B samples, which corresponds to 0.03x genome coverage. Following the clustering of the pooled reads and annotation of repeats represented by the largest clusters containing at least 0.01% of analyzed reads, proportions of reads in each cluster from 0B and +B samples were determined by parsing read names where genotypes of origin were encoded by specific tags. This allowed for identification of repeats enriched on B chromosomes as the clusters with increased proportions of reads from +B sample compared to 0B. Monomer lengths and consensus sequences of newly identified satellite repeats were determined using TAREAN (Novák et al., 2017). PCR primers were designed for consensus sequences of identified B-repeats (Suppl. Table 2 and 3). Control primers were designed for the *T. aestivum* 18S ribosomal RNA coding part of rDNA 25S-18S intergenic region *Eco*RI-*Bam*HI fragment (GenBank accession X07841.1). 0B/+B genomic DNA was used as template and PCR products were cloned using CloneJET PCR Cloning Kit (Thermo Fisher Scientific, Germany) and verified by sequencing (GeneBank, AesTR-126 MK283665,AesTR-148 MK283666, AesTR-183 MK283667, AesTR-205 MK283668).

### Molecular phylogenetic analysis of AesTR-183-like repeats

AesTR-183-like sequences were amplified by PCR from genomic DNA of *Ae. mutica*, *S. cereal*e and three *Ae. speltoides* accessions (Katzir, Technion, Ramat Hanadiv) all containing B chromosomes. Sequences were analyzed and assembled by Sequencher 5.4.6 (Gene Codes Corp., Ann Arbor, MI, USA). These sequences including a similar region of wheat chromosome 1B (accession LS992081.1, position 432930807-432930254) were aligned with ClustalW as implemented in Unipro UGENE v1.31.0 (Okonechnikov et al., 2012). When necessary, alignments were edited manually. jModelTest2 (Darriba et al., 2012) was used to estimate best-fit nucleotide substitution model resulting in selection of HKY + G out of 88 tested models according to the Bayesian information criterion (BIC). Phylogenetic analysis was performed in MrBayes v3.2.6 (Ronquist and Huelsenbeck, 2003) by running two parallel Metropolis-coupled Markov chain Monte Carlo analyses with four chains each for 2 million generations. Tree sampling was conducted every 500 generations. The consensus tree was generated after removal of 25% of samples from the beginning of the chain as burn-in. The final tree was visualized with FigTree v1.4.3 (http://tree.bio.ed.ac.uk).

### Flow sorting of sperm and vegetative nuclei

Mature pollen grains from anthers at dehiscence stage were collected and the corresponding nuclei were isolated by applying the filter bursting method (Kron and Husband, 2012) using CellTrics disposable filters of 100 μm and 20 μm (SYSMEX). The resulting nuclei suspension was stained with propidium iodide (50 μg/ml, PI) and analyzed on a BD Influx (Becton Dickinson) by blotting the PI fluorescence against the forward scatter signal. Based on differences in the DNA content and nuclear size, populations of vegetative and sperm nuclei without and with Bs could be differentiated. The individual fractions were flow-sorted into Eppendorf tubes and fixed with 4% formaldehyde in 5% sucrose buffer (10 mM Tris, 50 mM KCl, 2 mM MgCl_2_, 5% sucrose, 0.05% Tween, pH 7.5) for 10 min. Subsequently, 15-20 μl of the nuclei suspensions were applied on microscopic slides and dried overnight before performing immunostaining and FISH.

### Preparation of tissue sections and chromosome spreads

For the preparation of tissue sections, post-meiotic anthers (5 - 5.5 mm in length) were fixed in 4% paraformaldehyde prepared in 1× MTSB buffer as described in (Houben et al., 2011). In brief, for embedding, the anthers were infiltrated first with a graded series of PEG-wax in absolute ethanol solutions (1/2, 1/1, 2/1, (v/v)), 12 h for each at 37°C. Finally, anthers were infiltrated with pure PEG-wax for 12 h at 37°C and transferred into casting moulds. 10 μm tissue sections were cut on histological microtome Leica RM 2265 (Leica Biosystems) and placed on poly-L-lysine coated microscopic slides. The preparation of meiotic chromosomes was performed according (Badaeva et al., 2017). Meiotic anthers (~1.5 mm in length) were fixed in 3:1 (ethanol: acetic acid, v/v) for one hour. Meiocytes were squeezed out of the anthers on a slide in 7 μl 45% acetic acid with the help of a needle and squashed between slide and coverslip. The coverslips of liquid nitrogen frozen slides were removed and slides were kept in 1×PBS at 4°C for immunostaining and FISH. The preparation of mitotic chromosomes from the apical meristem of three days old seedlings was performed according to Aliyeva-Schnorr et al. (2015) by dropping the cell suspension on the slides. The slides were stored until use in 96% ethanol at 4℃.

### Immunostaining and FISH

Slides carrying the anther tissue sections were washed in 96% ethanol twice for 10 min, in 90% ethanol twice for 10 min, and finally in dH_2_O twice for 5 min. After an 800 Watt microwave treatment of the slides for 60 seconds (Chelysheva et al., 2010) in 10 mM sodium citrate buffer, slides were immediately transferred into a 1×MTSB for 5 min. Mouse anti-α tubulin (Sigma, cat. no. T 9026, diluted at 1:200), rabbit anti-grass CENH3 (Sanei et al., 2011) (diluted at 1:3000) and rabbit anti-histone H3K27me3 (Millipore, #07-449, diluted at 1:100) were used as primary antibodies. Anti-mouse Alexa 488 (Molecular Probes, cat. no. #A11001, diluted in 1:200) and anti-rabbit Cy3 (Jackson/ Dianova, cat. no. 111-165-144, diluted in 1:300) were used as secondary antibodies. To combine FISH and immunostaining, slides were rinsed with dH_2_O and placed in 2×SSC for 5 min before FISH. FISH was performed after post-fixation of specimens in 3:1 fixative for 10 min, washes in 2 × SSC twice and final fixation in 3% paraformaldehyde. 50 μl hybridization mixture per slide were used and the denaturing of tissue sections was done for 5 min on a hot plate at 80°C. For multiple rounds of FISH, FISH signal stripping was performed as described by (Heslop-Harrison et al., 1992; Shibata et al., 2009). In brief, after removing the coverslips the slides were rinsed in 0.1 × SSC at room temperature for 2 × 5 min. Next, slides were incubated in shaking jars at 42 °C for 30 min in each solution: once in 0.1 × SSC, twice in probe stripping solution [0.05% (v/v) Tween-20, 50% (v/v) formamide in 0.1 × SSC] and once in 4T solution [0.05% (v/v) Tween-20 in 4 × SSC]. Finally, specimens were fixed in 4% formaldehyde for 10 min, dehydrated in an ethanol series and air-dried.

### Generation of FISH probes

The wheat (peri)centromere-specific probes 192 bp (Ito et al., 2004), 6C6-3&4 (Zhang et al., 2004), CCS1 (Aragón-Alcaide et al., 1996), centromere satellite repeat (CenSR) (Li et al., 2013), CRW2 (Liu et al., 2008), pBs301(Cheng and Murata, 2003), RCS2 (Liu et al., 2008), Quinta-LTR (Li et al., 2013), TaiII (Kishii et al., 2001) and Weg 1-LTR (Li et al., 2013) were generated by PCR (Suppl. Table 2). Nine of the eleven probes are part of a (peri)centromeric Ty3/gypsy-type retrotransposon. The corresponding positions along the retrotransposon are shown in Suppl. Figure 7. Plant genomic DNA was extracted by the CTAB method (Murray and Thompson, 1980). The 25 μl PCR reactions contained following components: 2.5 μl 10×PCR buffer (20 mM MgCl_2_), 1.5 μl dNTP mix (10 mM), 1 μl of each primer (10 mM), 0.2 μl *Taq* (1U) polymerase (Quiagen), 50 ng gDNA and dH_2_O. Purified PCR products as well as plasmid clones of the newly identified repeats, the rye B-specific repeat D1100 (Sandery et al., 1990) and 5S rDNA clone pTa794 (Gerlach and Dyer, 1980) were directly labelled with dUTP-ATTO-647, dUTP-ATTO-550 or dUTP-ATTO-488 using a nick translation labelling kit (Jena Bioscience, http://www.jenabioscience.com).

### Microscopy and image analysis

The specimens were analyzed using an Olympus BX61 microscope equipped with an ORCA-ER CCD camera (Hamamatsu, Japan). Deconvolution microscopy was employed to remove out-of-focus information. Maximum intensity projections were processed with the program software CellSens dimension (Olympus). All images were collected in grey scale and pseudo-coloured with Adobe Photoshop CS (Adobe). The ImageJ (https://imagej.nih.gov/ij/download.html) particle analyzis tool was used to measure the size of CENH3 signals. Graphs showing the CENH3 signal size and the length of spindel microtubules were prepared with the Rstudio ggplot2 package.

## Supplemental material

The following materials are available in the online version of this article.

## Additional figures

**Figure S1.** PCR amplification of AesTR-126, AesTR-148, AesTR-183 and AesTR-205 repeats from *Ae. speltoides* genomic DNA of 0B and +B plants. **Figure S2.** Distribution of the B-specific probes AesTR-183 (magenta) and AesTR-205 (green) on the two B chromosome variants of *Ae. speltoides.* **Figure S3.** Phylogeny of AesTR-183-like sequence. **Figure S4.** Flow cytometry efficiently showed B chromosome accumulation in sperm nuclei of +B plants compared with a 0B plant. **Figure S5.** Scheme of the B chromosome transmission process in the male gametogenesis of *Ae. speltoides* exemplified for plants with 1B, 2B and 3B chromosomes. **Figure S6.** Sequence composition of *Ae. speltoides* A and B (peri)centromeres. **Figure S7.** Schemata of a Ty3/gypsy retrotransposon showing the positions of the used (peri)centromeric FISH probes.

## Additional Table

**Table S1.** Repeat composition of *Ae. speltoides* 0B and +B genotypes estimated from clustering analysis of Illumina reads. **Table S2.** PCR primers used for amplification of AesTR-126, AesTR-148, AesTR-183 and AesTR-205 repeats, control primers for 18S rDNA and primers for (peri)centromeric sequences. **Table S3.** Consensus monomer sequences of AesTR-126, AesTR-148, AesTR-183 and AesTR-205 determined by TAREAN analysis and used to design the primers.

## Acknowledgements

We thank Olga Raskina (Institute of Evolution, University of Haifa, Israel) and John Harper (Aberystwyth University, UK) for providing *Ae. speltoides* and *L. perenne*, respectively. We thank Katrin Kumke and Oda Weiss for technical assistance, as well as Nadine Bernhardt, Frank Blattner and Ingo Schubert for insightful discussions. This work was supported by the China Scholarship Council (Grant No. 201606910015) (to DW), the Czech Academy of Sciences (Grant No. RVO:60077344) (to PN, JM) and the Deutsche Forschungsgemeinschaft DFG (Grant No. HO1779/26-1) (to AH).

The authors declare that they have no competing interests.

## Author contributions

DW, AR, JF, YZ and AH designed the study, interpreted the data and wrote the manuscript. MV, PN, JM performed the bioinformatics analysis. JF conducted the flow cytometry. All authors read and approved the final manuscript.

## List of abbreviations

A: A chromosome
B: B chromosome
BAD: Background aggregate and debris percentage
CENH3: Centromere histone H3
ChIP: Chromatin immunoprecipitation
CV: Coefficient of variation
DAPI: 4′,6-diamidino-2-phenylindole
DNA: Deoxyribonucleic acid
FISH: Fluorescence in situ hybridization
FSC: Forward scatter signal
Gen: Generative nucleus
H3K27me3: Trimethylated histone H3 at lysine 27
Micro: Micronucleus
MTSB: Microtubule stabilization buffer
PCR: Polymerase chain reaction
PI: Propidium iodide
RNA: Ribonucleic acid
Sp: Sperm nucleus
SSC: Saline sodium citrate buffer
Veg: Vegetative nucleus

